# Computational model for Rumination as Self-Supervised Learning in a World Model

**DOI:** 10.64898/2026.11.22.768285

**Authors:** Kengo Inutsuka, Masato Tsutsumi, Iori Higashino, Yasushi Okochi, Takatoshi Hikida, Takahiro A. Kato, Honda Naoki

**Author notes:** These authors contributed equally to this work (Kengo Inutsuka, Masato Tsutsumi).

## Abstract

Rumination is a cognitive process characterized by repetitive negative thoughts that persist even in the absence of external threats. Although excessive rumination is associated with stress-related mental disorders, the computational mechanisms underlying its persistence and its impact on threat perception remain unclear. Here, we propose a computational framework in which rumination emerges from recursive learning within a world model. Inspired by the social defeat stress (SDS) paradigm in mice, our model assumes that direct experiences of social defeat first establish beliefs about the threat posed by an aggressor. These beliefs subsequently generate internally simulated experiences, which are repeatedly used to update the beliefs themselves. We investigated how this recursive process reshapes threat perception by allowing the rumination gain, a learning rate controlling internal belief updating, to take both positive and negative values. Through numerical simulations and mathematical analysis, we found that positive gains did not simply reinforce acquired threat beliefs, while negative gains did not simply erase them. Instead, both positive and negative gains could generate markedly divergent threat perceptions, with some simulated individuals perceiving severe threats and others perceiving almost none, despite identical initial beliefs and external experiences. These outcomes qualitatively resemble the coexistence of stress-susceptible and resilient mice following SDS. Our findings suggest that rumination is not merely a repetition of past experiences but a self-referential learning process that can transform identical stressful experiences into fundamentally different perceptions of threat.

## 2 INTRODUCTION

Rumination is a cognitive process characterized by repetitive and persistent attention to negative thoughts and experiences [1]. Rather than representing a transient emotional response, rumination often emerges as a self-sustaining mental process that continues to dominate internal cognition even when the eliciting stressor is no longer present [2, 3]. Excessive rumination is associated with persistent affective disturbances and reduced cognitive flexibility [2, 4]. However, despite existing computational accounts [5, 6], the mechanisms that sustain rumination over time and render it self-reinforcing remain unclear. Such persistence motivates examining how internally generated experiences may reshape beliefs about threat beyond direct exposure to a stressor.

Here, we use the term world model for an internal probabilistic representation of how the environment evolves and generates observations. This framework allows us to examine how internally generated observations can sustain repeated learning. Humans and animals are thought to rely not only on immediate sensory input but also on internal models learned from past experience to anticipate future outcomes and social interactions [7–10]. A key property of world models is that they can generate predictions and simulated experiences from learned representations without requiring a new external observation at each step. While this capability supports adaptive learning and decision-making [11], it also entails an inherent vulnerability: under conditions of strong or prolonged stress, internally generated predictions and evaluations may become excessively repeated and reinforced. Rumination can be understood as a state where learning and updating within the world model become biased in a specific direction. This provides a mechanistic explanation for the persistent cognitive changes observed after stress exposure.

The tendency for rumination to emerge after stress exposure has been reported in human studies [3, 12, 13], motivating the development of experimental models to elucidate its neural and behavioral underpinnings. Among these, Social Defeat Stress (SDS) involves repeated experiences of social subordination and is widely adopted as a standardized animal model reliably inducing persistent social avoidance behavior and emotional changes [14, 15]. The defining feature of SDS lies in its temporal structure: brief, intense stressors are repeatedly administered, followed by prolonged psychological stress [15]. This qualitative and temporal asymmetry creates conditions for altered internal cognitive processes.

A prominent and unresolved feature of the SDS paradigm is the marked inter-individual variability in behavioral outcomes. Even when exposed to identical stress protocols, some animals exhibit robust and persistent fear responses and social avoidance, while others display relatively normal behavior and resilience to stressors [14]. This dichotomy of susceptibility-resilience shows high reproducibility across laboratories, but the mechanism by which the same stress experience generates different internal cognitive trajectories over time remains unclear.

From a world model perspective, SDS can be viewed as an environment where access to diverse novel external information is restricted, and negative predictions/evaluations are repeatedly activated. Under such conditions, internally generated predictions and memories tend to dominate the learning process, causing world model updates to skew toward self-reinforcing dynamics. Despite its widespread use, SDS has been scarcely examined from a computational perspective that explicitly models the internal cognitive dynamics and recursive learning induced by stress protocols.

Here, we develop a computational model of rumination as recursive learning within a world model. In the model, threat beliefs shaped by social defeat guide the generation of simulated experiences, which are then used to update those same beliefs. We investigate how rumination reshapes threat beliefs by varying the rumination gain, a learning rate controlling belief updates during internal simulation. Through numerical simulations and mathematical analysis, we show that rumination can generate divergent threat perceptions irrespective of the sign of the rumination gain, even from identical initial beliefs and experiences. These findings are qualitatively consistent with the coexistence of stress-susceptible and resilient mice following SDS. Our findings suggest that rumination is not merely a repetition of past experiences but a process that can fundamentally reshape how those experiences are perceived.

## 3 MODELS

### 3. Rumination as recursive Bayesian updating

In this study, we define a world model as an internal probabilistic representation of how the environment evolves and generates sensory inputs. In the Bayesian brain framework, perception involves beliefs about the unobserved causes of sensory input ([16]). We denote these latent causes by *Z* and observations by *O*. Prior beliefs and an observation model *P* (*O*|*Z*) together specify how observations are expected to arise from latent causes (Fig. 1a).

**Fig. 1.**
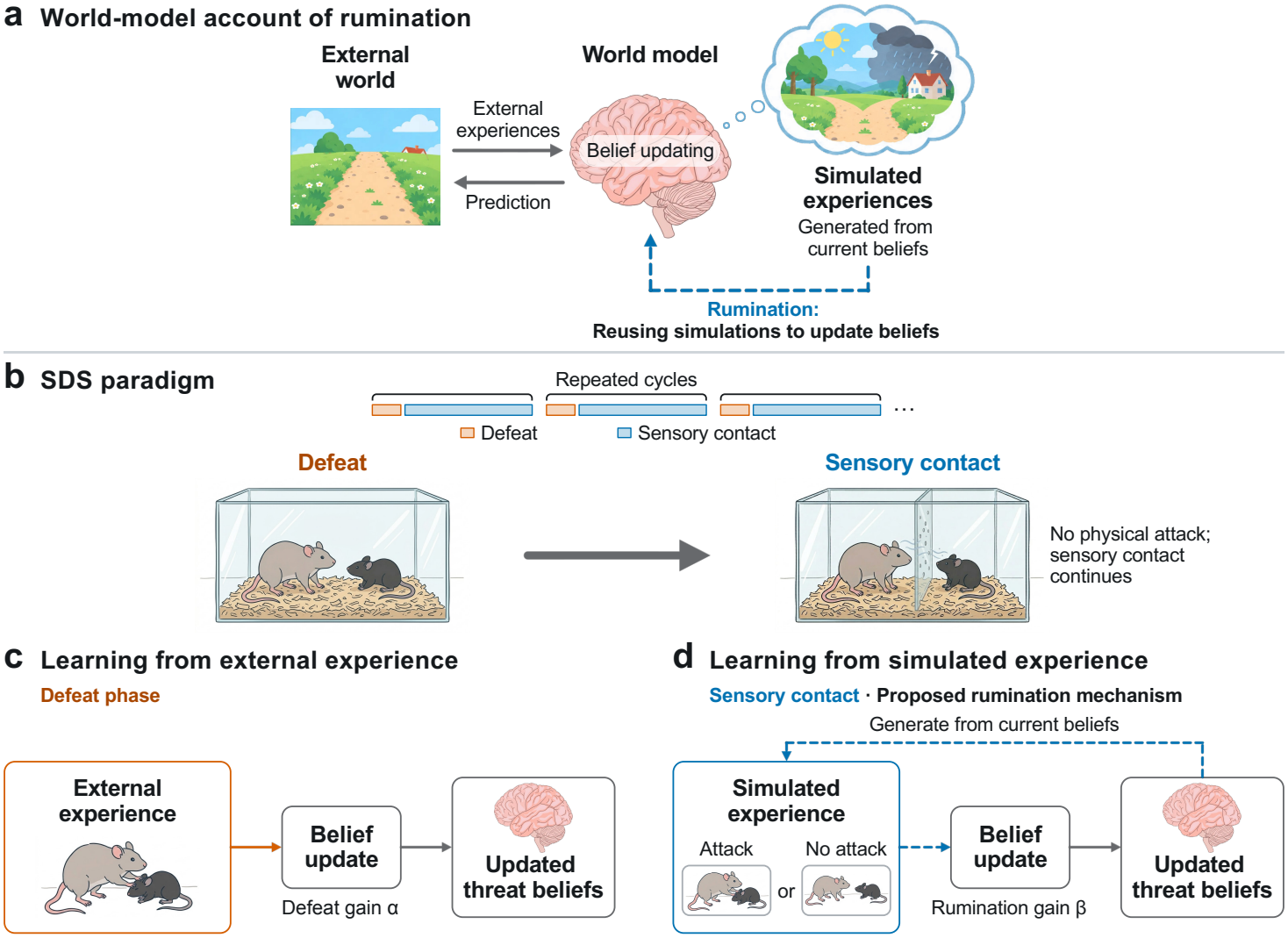
A world-model framework for rumination under social defeat stress. **a**, Conceptual relationship between world models and rumination. External experiences update beliefs within a world model, which supports predictions and generates simulated experiences. Rumination is modeled as repeated belief updating using these internally generated experiences. **b**, Social defeat stress (SDS) paradigm. Brief defeat phases (orange) alternate with prolonged sensory-contact phases (blue), during which the mouse receives sensory cues from the aggressor while a perforated divider prevents physical attacks. **c**,**d**, The proposed model of rumination is illustrated using the defeat (c) and sensory-contact (d) phases of SDS. **c**, Attacks provide external observations for updating threat beliefs (orange solid arrow). The defeat gain *α* scales updates to the mean threat belief driven by external attack observations. **d**, Simulated observations of attack or no attack are generated from current threat beliefs and reused to update them in a recursive loop (blue dashed arrows). The rumination gain *β* controls the direction and magnitude of updates to the mean threat belief.

Bayesian inference combines prior expectations with the evidence provided by an observation to obtain an updated, or posterior, belief. At step *t*, the posterior belief after incorporating an observation *O*_*t*_ is

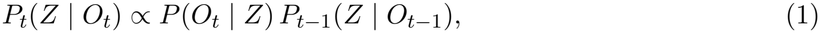

where *P*_*t−*1_(*Z* |*O*_*t−*1_) is the belief before incorporating *O*_*t*_. The observation changes the relative plausibility of possible latent causes according to how well each could account for it. Thus, *P*_*t*_(*Z* |*O*_*t*_) represents the belief after incorporating the observation, while retaining the influence of prior expectations.

Within this framework, we define rumination as a recursive inference process driven by internally generated observations (Fig. 1a). At each step, a latent cause is sampled from the current posterior belief, and an observation is generated from the observation model:

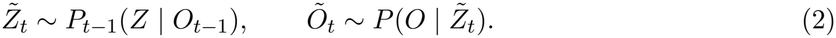

The internally generated observation *Õ*_*t*_ is then treated as new evidence and used to update the belief:

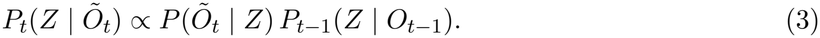

This process is repeated, creating a recursive loop in which internally generated observations continue to drive learning even when little or no new external information is available. The observation model remains unchanged; the source of the observations used for learning shifts from external events to internal generation.

### 3.2 Modeling rumination under social defeat stress

To separate the contributions of direct attacks and subsequent internal learning to threat beliefs, we model each session of social defeat stress (SDS) as two consecutive phases: a defeat phase in which a mouse is attacked by an aggressor, followed by a sensory-contact phase in which a perforated divider prevents physical interaction while allowing sensory contact (Fig. 1b).

The latent variable *Z* represents an unobserved threat level that determines the aggressor’s likelihood of attacking. At step *t*, the mouse’s belief about its possible values *z* is represented by a Gaussian recognition distribution,

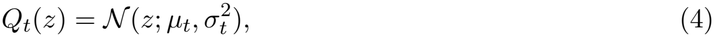

where *µ*_*t*_ is the mean threat belief and 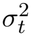 is the recognition variance, representing uncertainty about latent threat.

The world model has two components. The system model allows latent threat to change over time, adding system noise with variance 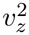 to the belief variance at each inference step. The observation model maps a latent threat to the probability of an attack through the sigmoid function 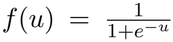. Together, they form the state-space model underlying the recognition updates described in Methods, Section 6.1.

#### 3.2.1 Phase 1: Defeat (external evidence-driven learning)

During the defeat phase, the mouse receives an external observation *o*_*t*_, a binary realization of *O*_*t*_, where *o*_*t*_ = 1 indicates an attack and *o*_*t*_ = 0 indicates no attack. In this phase, each observation is an attack (*o*_*t*_ = 1; Fig. 1c). The mean and precision of the threat belief are updated with defeat gain *α* according to

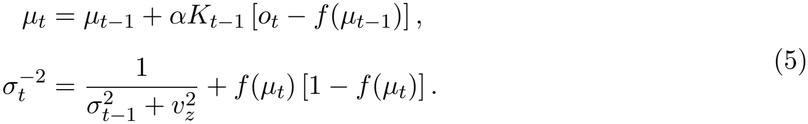

Here, *f* (*µ*_*t−*1_) is the predicted probability of an attack, and the term in brackets is the prediction error. The predictive variance 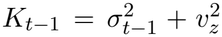 combines belief uncertainty and system noise, whereas *α* scales the mean update. The first term in the precision update incorporates uncertainty from the preceding step and system noise, while the second is the contribution of the observation model evaluated at the updated mean *µ*_*t*_. These updates establish the belief state entering sensory contact (Methods, Sections 6.1.1 and 6.1.2).

#### 3.2.2 Phase 2: Sensory contact (self-generated learning; rumination)

During sensory contact, physical attacks cease while sensory exposure to the aggressor continues (Fig. 1d). We hypothesize that rumination sustains internal learning during this interval. At each step, the mouse samples a threat level from its current belief and generates a virtual observation through the observation model:

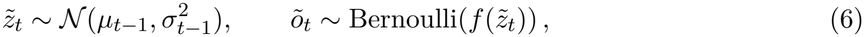

where *õ*_*t*_ = 1 represents a virtual attack and *õ*_*t*_ = 0 represents its absence. The generated observation is then used to update the belief, with rumination gain *β*:

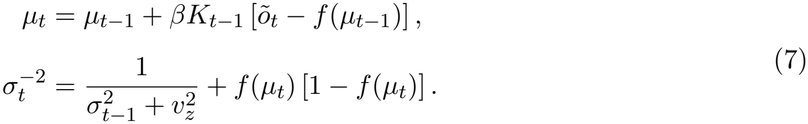

Repeating this process implements the recursive loop introduced above.

#### 3.2.3 Interpretation of *β*

We implement recursive learning with a recognition rule in a Bayesian state-space framework. The sign of *β* determines the direction of belief updating, while its magnitude controls the strength of the update. Intuitively, positive *β* may reinforce threat beliefs and amplify perceived threat beyond actual experiences, whereas negative *β* may act like forgetting, weakening acquired threat beliefs and alleviating rumination. At *β* = 0, the belief mean remains unchanged, although uncertainty can still evolve. We therefore investigate whether these intuitive expectations hold when internally generated experiences are repeatedly used to update beliefs.

## 4 RESULTS

### 4.1 Increasing positive rumination gain amplifies fl uctuations an d po larizes threat perception

To investigate how threat beliefs evolve during SDS, we simulated repeated sessions, each comprising a defeat phase followed by a sensory-contact phase. We quantified p erceived threat a s *q* _*t*_ = *f* (*µ*_*t*_), where *µ*_*t*_ is the mean threat belief and larger *q*_*t*_ indicates a higher perceived likelihood of attack. As an illustrative example, Fig. 2a shows the first t wo s essions o f a s ingle s imulation r un with positive defeat and rumination gains. During defeat, repeated experiences of attack consistently increased perceived threat. In contrast, during sensory contact, rumination driven by stochastically generated experiences caused irregular fluctuations in perceived t hreat. These contrasting dynamics suggest that internally generated experiences introduce variability into threat perception even in the absence of further physical attacks.

**Fig. 2.**
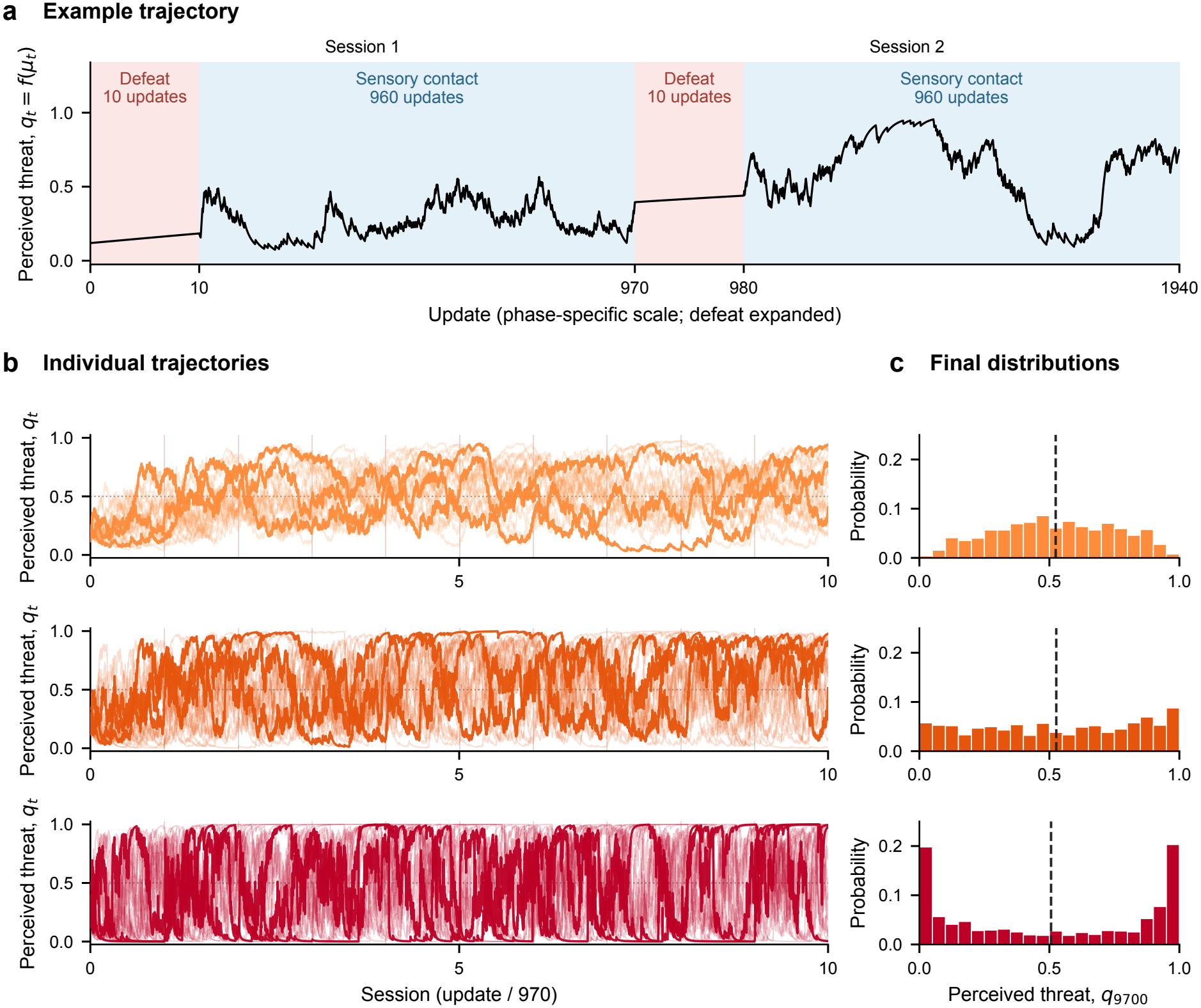
Trajectories and final distributions of perceived threat with positive rumination gains. Simulations use defeat gain *α* = 0.136 over 10 sessions, with 10 defeat updates followed by 960 sensory-contact updates per session. All other parameters and the initial conditions are the same across simulations (Methods, Section 6.2). Perceived threat is quantified as *q*_*t*_ = *f* (*µ*_*t*_), where *µ*_*t*_ is the mean threat belief and *f* is the sigmoid function. Light orange, dark orange, and red indicate *β* = 0.48, 1.04, and 1.96, respectively, in panels **b** and **c. a**, First two sessions of the first simulation run at *β* = 1.04, also included in **b**. Red and blue shading indicate defeat and sensory contact, respectively. The horizontal axis is linear within each phase, with defeat expanded for visibility (display-width ratio, 1:4); tick labels show cumulative updates. **b**, Individual trajectories from the first 20 of the 1, 024 simulation runs per gain, with the first three highlighted. The horizontal axis is linear in update count, with 970 updates per session; narrow red bands indicate defeat phases. Horizontal dotted lines mark *q*_*t*_ = 0.5. **c**, Final distributions of perceived threat (*q*_*T*_ at *T* = 9700) across all 1,024 runs per gain. Bars show fractions of runs in bins of width 0.05 on [0, 1]; vertical dashed lines indicate across-run means. In order of increasing *β*, across-run means (sample SDs) are 0.524 (0.229), 0.526 (0.307), and 0.506 (0.387).

We next examined how the rumination gain *β* affects the dynamics of perceived threat. Because simulated experiences are stochastically generated from current beliefs, each simulation run produces a different trajectory. We performed multiple simulation runs for each of three positive gains: small, intermediate, and large (Fig. 2b). At the smallest gain, perceived threat fluctuated substantially but remained mostly at intermediate values around 0.5. As *β* increased, fluctuations became more pronounced, with perceived threat spending longer periods near either extreme, 0 or 1. Some trajectories also switched between these opposing threat perceptions over time.

To examine how these dynamics shape the final outcomes, we analyzed the distribution of perceived threat at the end of the simulations (Fig. 2c). At the smallest gain, the distribution was unimodal and centered near 0.5. As *β* increased, the distribution became progressively broader, eventually exhibiting a bimodal, U-shaped distribution at the largest gain, with marked accumulation near 0 and 1. Notably, the mean across simulation runs remained close to 0.5 for all three gains, despite the increasing divergence among individual runs. Thus, contrary to the intuitive expectation that repeated rumination would reinforce acquired threat beliefs, stronger positive rumination gains produced highly fluctuating threat perceptions and increasingly polarized outcomes, ranging from almost no perceived threat to nearly certain attack.

### 4.2 Negative rumination gains polarize threat perception and shift outcomes toward high threat

We next examined how negative rumination gains (*β <* 0) affect perceived threat. We performed simulations for three negative gains with small, intermediate, and large magnitudes (*β* = −0.48, −1.04, and −1.96, respectively; Fig. 3a). At the smallest gain magnitude, perceived threat increased during defeat phases but decreased toward 0 during sensory contact in most trajectories. However, some trajectories gradually approached 1 and remained near 1 for the rest of the simulation. At intermediate and large gain magnitudes, transitions toward either 0 or 1 became much faster. In many trajectories, perceived threat rapidly approached 1 during sensory contact immediately after a defeat phase. This tendency became more pronounced as *β* became more negative.

**Fig. 3.**
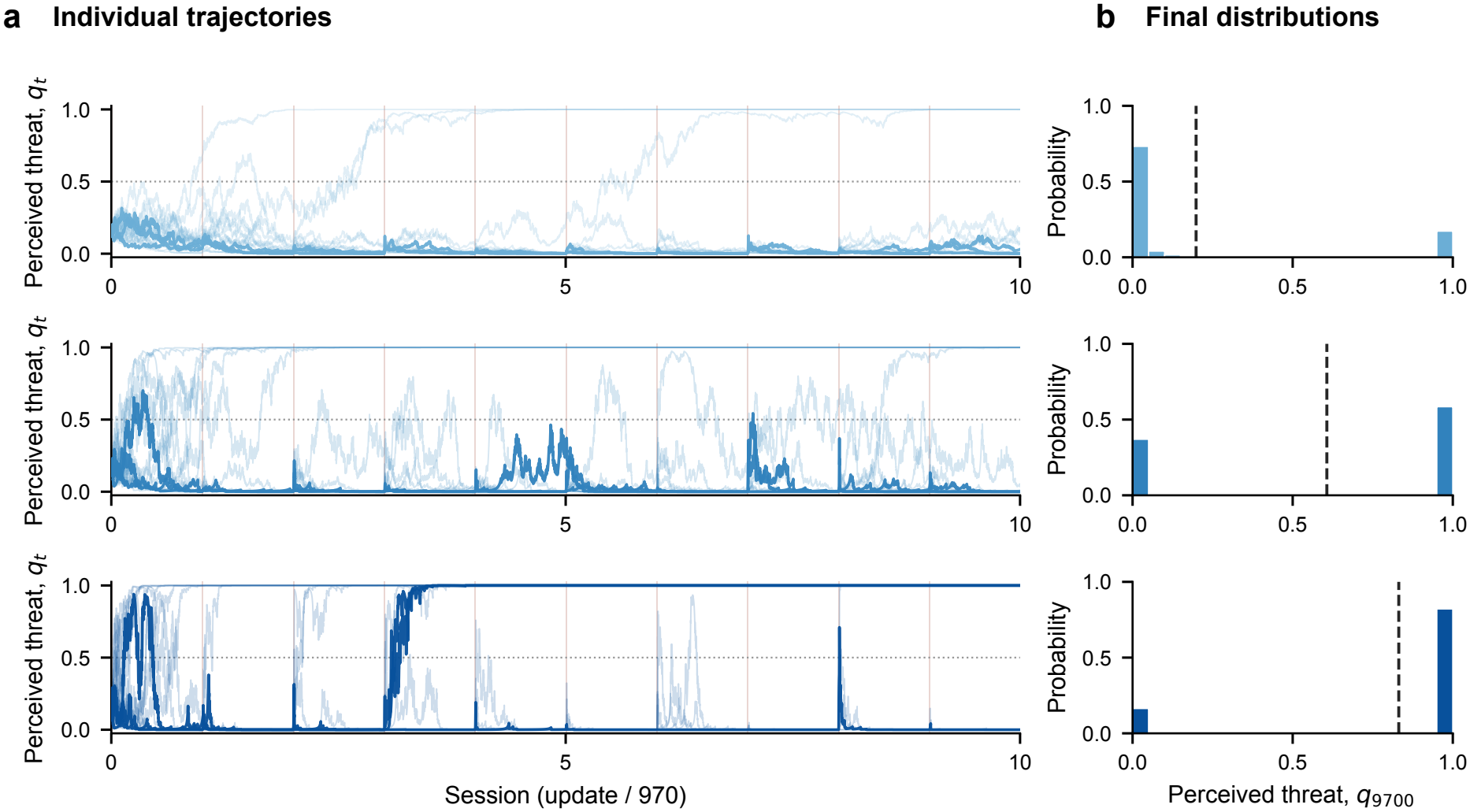
Trajectories and final distributions of perceived threat with negative rumination gains. Simulations use the same protocol, initial conditions, and parameter values as in Fig. 2, except for the rumination gain *β*. The defeat gain is *α* = 0.136. Perceived threat is quantified as *q*_*t*_ = *f* (*µ*_*t*_), where *µ*_*t*_ is the mean threat belief and *f* is the sigmoid function. Light, medium, and dark blue indicate *β* = *−*0.48, *−*1.04, and *−*1.96, respectively. **a**, Individual trajectories from the first 20 of the 1, 024 simulation runs per gain, with the first three highlighted. The horizontal axis is linear in update count, with 970 updates per session; narrow red bands indicate defeat phases. Horizontal dotted lines mark *q*_*t*_ = 0.5. **b**, Final distributions of perceived threat (*q*_*T*_ at *T* = 9700) across all 1,024 runs per gain. Bars show fractions of runs in bins of width 0.05 on [0, 1]; vertical dashed lines indicate across-run means. In order of increasing gain magnitude, across-run means (sample SDs) are 0.198 (0.378), 0.607 (0.481), and 0.831 (0.372).

The final distributions of perceived threat were bimodal across all negative gains examined, with outcomes concentrated near 0 and 1 (Fig. 3b). At the smallest gain magnitude, the distribution was biased toward 0. As *β* became more negative, the proportion of outcomes near 0 decreased, while the proportion near 1 increased. Thus, negative rumination gains did not uniformly reduce perceived threat: more negative *β* values favored high-threat outcomes. Together with the results for positive gains, these findings show that recursive belief updating during rumination can polarize threat perception under both positive and negative rumination gains.

### 4.3 Stability and stochastic fluctuations explain gain-dependent threat dynamics

The simulations showed that both positive and negative rumination gains could polarize threat perception, but with distinct temporal dynamics. At large positive gains, some trajectories repeatedly switched between perceived threat near 0 and near 1, whereas negative gains produced trajectories that remained near either 0 or 1 (Figs. 2b and 3a). We then examined how perceived threat at the end of the simulations varied across a broad range of the defeat gain *α* and rumination gain *β* (Fig. 4a). As *β* increased within the positive range, the mean perceived threat across simulation runs approached 0.5, while the standard deviation generally increased. For *β <* 0, the simulation outcomes were dominated by either low or high perceived threat, depending on the parameter values. Along the white dotted line in Fig. 4a, the outcomes shifted toward higher perceived threat as *β* became more negative, consistent with Fig. 3b. Within the negative-gain region, the standard deviation was highest where outcomes were roughly evenly divided between values near 0 and 1.

**Fig. 4.**
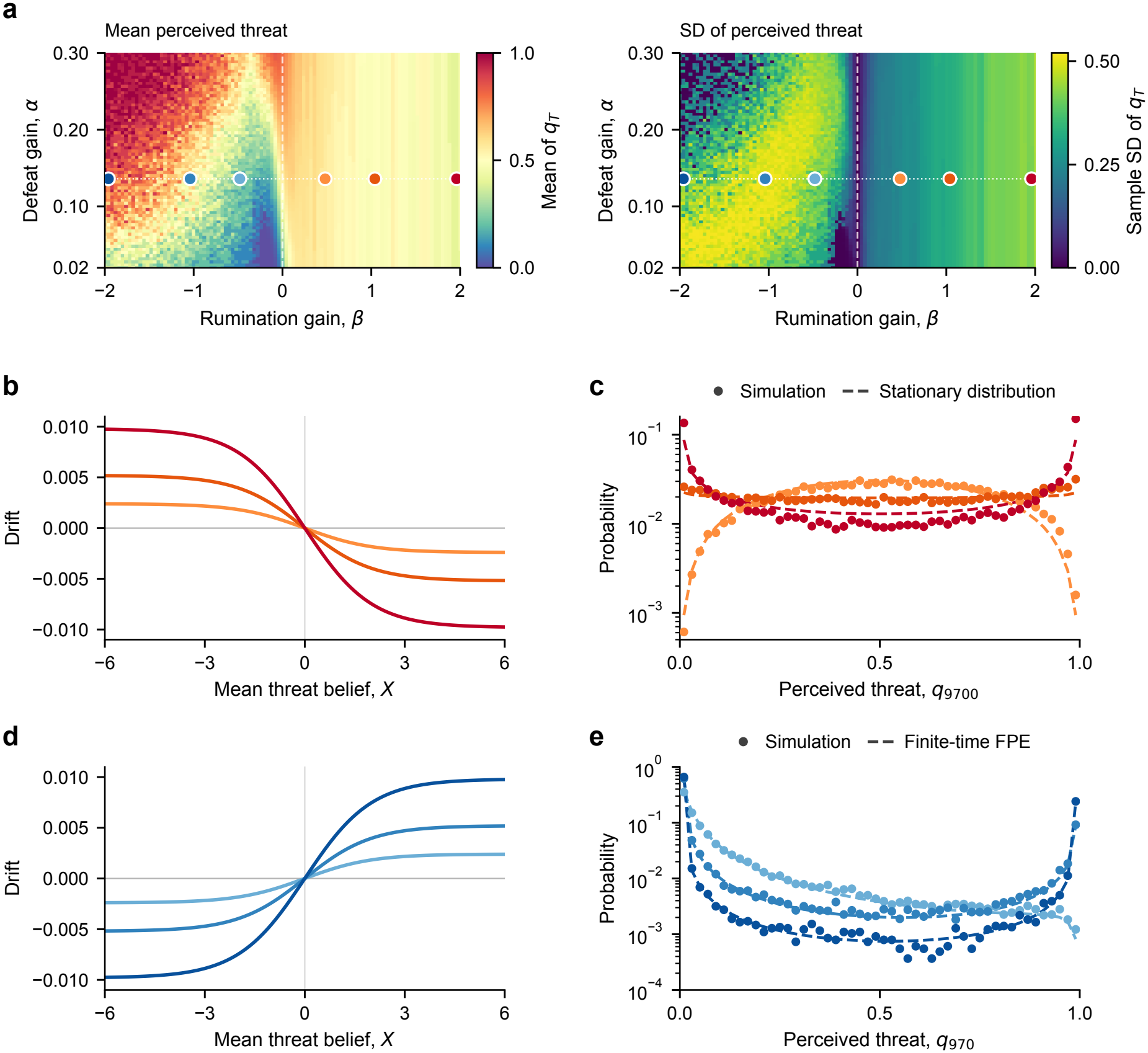
Gain-dependent perceived threat statistics and continuous-time approximation of sensory-contact dynamics. Perceived threat is quantified as *q*_*t*_ = *f* (*µ*_*t*_), where *µ*_*t*_ is the mean threat belief and *f* is the sigmoid function (see Methods, Section 6.2 for simulation protocol and initial conditions). Light orange, dark orange, and red indicate *β* = 0.48, 1.04, and 1.96, respectively; light, medium, and dark blue indicate *β* = *−*0.48, *−*1.04, and *−*1.96, respectively. **a**,, Mean (left) and sample SD (right) of final perceived threat (*q*_*T*_ at *T* = 9700) across 20 simulation runs per parameter pair for *α ∈* [0.02, 0.30] and *β ∈* [*−*2, 2]. Circles mark the six gain conditions used in Figs. 2 and 3 along *α* = 0.136 (horizontal dotted line); the vertical dashed line marks *β* = 0. **b**,,**d**,, Theoretical drift of the continuous-time mean threat belief *X* (Eq. (8), with 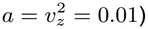, pointing toward *X* = 0 for positive gains (**b**,) and away from *X* = 0 for negative gains (**d**,). **c**,,**e**,, Perceived threat distributions (logarithmic vertical axes; bin width 0.02 on [0, 1]) comparing simulations (circles; 16,384 runs per gain at *α* = 0.136) with continuous-time analytical and numerical predictions (dashed curves). For positive gains (**c**,), simulated distributions at *T* = 9700 (after 10 sessions) are compared with the stationary Beta(1/*β*, 1/*β*) distribution (Eq. (33)). For negative gains (**e**,), simulated distributions at *T* = 970 (after the first session of 10 defeat and 960 sensory-contact updates) are compared with finite-time Fokker--Planck equation (FPE) solutions initialized at the post-defeat mean threat belief (Methods, Section 6.4.3).

To understand how these two distinct temporal dynamics lead to polarized threat perception, we approximated belief updating from simulated experiences during sensory contact (Eq. (13) with *η*_*t*_ = *β*) as a continuous process. Assuming that the recognition variance is small and follows its local equilibrium value at the current belief mean, we applied a diffusion approximation to obtain the following stochastic differential equation for the belief mean (Methods, Section 6.4):

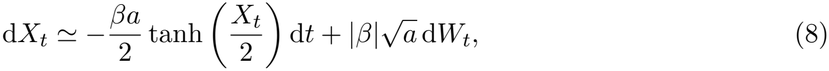

where *X*_*t*_ is the continuous-time counterpart of the mean threat belief 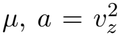 is system noise with variance, and *W*_*t*_ is a standard Wiener process. The first term represents the drift, whereas the second represents diffusion arising from stochastic fluctuations. The drift magnitude and noise amplitude both scale linearly with |*β*|.

For *β >* 0, the drift was directed toward *X* = 0, corresponding to a perceived threat of *q* = 0.5, making the center stable in the absence of fluctuations (Fig. 4b). We analytically derived the stationary distribution of *X*_*t*_ from Eq. (8). This distribution remained unimodal and symmetric around zero, but broadened as *β* increased. After transformation to perceived threat, *q* = *f* (*X*), the distribution changed from unimodal at small *β*, through a uniform distribution, to bimodal with peaks near 0 and 1 as *β* increased (Fig. 4c). Thus, although the distribution of *X* remained centered at zero, its broadening led to a concentration of perceived threat near the two extremes. At the same time, the restoring drift toward *X* = 0 allows trajectories to return from either extreme, consistent with the repeated switching observed in the simulations. This analytical result agreed qualitatively with the simulation results for positive *β* (Fig. 4c).

For *β <* 0, the drift was directed away from *X* = 0, corresponding to a perceived threat of *q* = 0.5, making the center unstable (Fig. 4d). Thus, the drift drives *X*_*t*_ away from the center, toward low or high perceived threat near 0 or 1. Once a trajectory approaches either extreme, the outward drift opposes its return toward the center, consistent with the prolonged residence near one extreme observed in the simulations. To assess the diffusion approximation, we numerically solved the Fokker–Planck equation associated with Eq. (8) to obtain the time-dependent distribution of *X*_*t*_ and transformed it into the distribution of perceived threat (Methods, Section 6.4.3). The resulting predictions agreed qualitatively with the simulation results for negative *β* (Fig. 4e). Although belief updates during sensory contact are symmetric around *µ* = 0, the initial belief and the preceding defeat can produce unequal proportions of outcomes near 0 and 1.

Together, these analyses show that threat perception can become polarized under both stable and unstable dynamics around the center, providing a theoretical explanation for the simulation results.

## 5 DISCUSSION

We modeled rumination as recursive learning from internally simulated experiences and showed that this process could polarize threat perception even from identical initial beliefs and external experiences. Across the three positive-gain conditions in Fig. 2, increasing the rumination gain *β* amplified fluctuations and shifted the final distribution of perceived threat from unimodal to bimodal, while the mean across simulation runs remained near 0.5. The negative gains examined also produced bimodal final distributions, with more negative values of *β* favoring high-threat outcomes (Fig. 3). At large positive gains, some trajectories repeatedly switched between low and high perceived threat, whereas negative gains produced prolonged residence near either extreme. For positive gains, the diffusion approximation showed that, despite a restoring drift toward *X* = 0, the stationary distribution of the belief mean *X* broadened as *β* increased while remaining unimodal. Applying the sigmoid transformation yielded a bimodal distribution of perceived threat at large gains. For negative gains, the drift was directed away from *X* = 0, driving trajectories toward low or high perceived threat and opposing their return toward the center (Fig. 4). Together, these findings show that threat perception can become polarized under both stable and unstable dynamics around the center.

Rumination can persist after a stressor has ended [3] and has been implicated in the link between stressful experiences and subsequent symptoms of depression and anxiety [12, 13]. These findings motivate examining how repeated internal processing reshapes threat beliefs acquired through stress. Previous computational accounts have examined recurrent negative-memory retrieval and its effects on task performance [5], persistent policy sampling that fails to resolve uncertainty [17], and prolonged sampling during hidden-state inference [6]. These accounts help explain how ruminative thought recurs or persists, but leave open how repeated learning from internally generated experiences changes the beliefs that generate subsequent experiences. Our model addresses this question by using simulated observations of attack or no attack to update the threat beliefs from which they were generated. The updated beliefs then shape subsequent simulated observations, forming a recursive learning process. Our findings show that this process can polarize threat perception rather than simply reinforce acquired threat beliefs, providing a computational account of how rumination can continue to reshape the cognitive consequences of stress after direct attacks have ceased.

In representative world-model approaches, such as those of Ha and Schmidhuber [18] and Hafner et al. [11], models learned from interaction with the environment generate simulated experiences for learning policies that maximize expected cumulative reward. Our model instead uses internally simulated experiences to update a recognition distribution over latent threat. The system and observation models are specified in advance and held fixed, while the mean and precision of the recognition distribution are updated through external and internally simulated experiences. This separates changes in threat beliefs from learning environmental dynamics or action policies.

The simulations identify a source of variability in threat perception that can complement explanations of stress susceptibility based on pre-existing individual differences. For example, prefrontal reactivity to an aggressor measured before chronic defeat predicts subsequent stress susceptibility [19]. Within each gain condition, simulation runs shared the same model parameters, initial beliefs, and external experiences. Because internally simulated experiences were generated stochastically, repeated belief updating produced different trajectories and final levels of perceived threat. This account is consistent with approaches in psychiatry that emphasize how mental states and vulnerability evolve over time [20, 21]. Our results motivate the hypothesis that rumination contributes to differences in stress susceptibility and resilience by generating divergent threat beliefs during sensory contact.

Distinguishing the temporal patterns that lead to polarized threat perception calls for longitudinal measurements during sensory contact. In the standardized SDS protocol, social interaction is assessed after the defeat series [15]; this endpoint alone does not capture how responses change between defeat phases. Willmore et al. [22] combined behavioral tracking with recordings of dopaminergic activity during defeat and brief sensory-contact phases before and after it, identifying behavioral and neural activity patterns associated with subsequent resilience. Extending these measurements over longer sensory-contact phases, together with a model linking threat beliefs to behavior and neural activity while accounting for the ongoing sensory input from the aggressor present in the standard SDS protocol, could allow individual time courses to be compared with the predicted patterns of fluctuation, switching, and prolonged residence near either extreme. Such analyses could examine how these temporal patterns relate to social interaction after the defeat series.

These longitudinal behavioral data could support joint inference of latent belief states and model parameters such as the rumination gain *β*. Inverse approaches have estimated animal behavioral strategies [23], the time-varying reward–curiosity balance [24], cognitive updating and optimism– pessimism bias [25], and latent world-model dynamics [26]. Applied to our model with appropriate attention to identifiability, such approaches could reconstruct the mean and variance of the recognition distribution over time and estimate the parameters governing belief updating. The inferred belief changes could then be related to behavioral phenotypes, such as social avoidance measured in social interaction tests, by incorporating a model of action selection. Active-inference modeling of social investigation and defensive behavior before and after social defeat [27] provides a starting point for this extension. The inferred belief trajectories could also be compared with neural activity, while individual differences in the estimated parameters and belief dynamics could be related to gene-expression profiles. Such comparisons could clarify how neural and molecular processes associated with stress susceptibility and resilience [22, 28] relate to recursive belief updating and its effects on threat perception.

## 6 MATERIALS AND METHODS

### 6.1 Discrete-time model and recognition updates

#### 6.1.1 World model and sequential inference

We model the mouse’s internal representation of threat using a state-space model. The latent variable *Z*_*t*_ represents the threat posed by the aggressor, with larger values corresponding to a higher probability of attack. The mouse assumes that latent threat evolves as a Gaussian random walk:

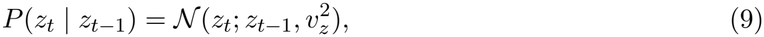

where 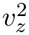 is the variance of latent change per update. This transition is an assumption of the mouse’s world model; the simulations do not generate an independent trajectory of the aggressor’s true threat.

The observation *o*_*t*_ represents either an attack (*o*_*t*_ = 1) or no attack (*o*_*t*_ = 0). We map latent threat to attack probability using the sigmoid function 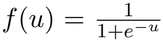. Given latent threat *z*_*t*_, the observation likelihood is

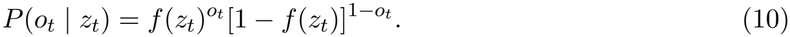

Thus, *f* (*z*_*t*_) gives the probability of an attack. The inference problem is to estimate latent threat sequentially from the observation history *o*_1:*t*_ = (*o*_1_, …, *o*_*t*_). The posterior distribution is updated according to

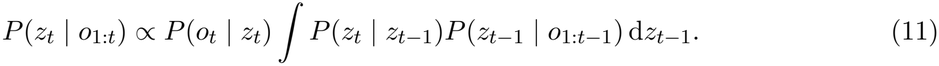

Because the Bernoulli observation model with a sigmoid link is not conjugate to a Gaussian distribution over latent threat, this update does not generally yield a Gaussian posterior. We therefore approximate the posterior at each step by a Gaussian recognition distribution:

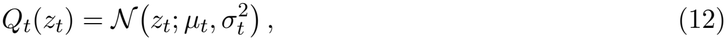

where *µ*_*t*_ is the mean threat belief and 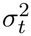 is its variance, the inverse of the recognition precision.

#### 6.1.2 Discrete update rule

Following the approximate variational free-energy formulation of Konaka and Honda [24], we update the mean *µ*_*t*_ and precision 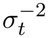 of the recognition distribution as follows:

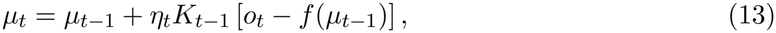

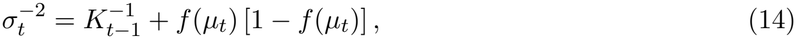

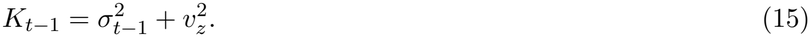

Here, *o*_*t*_ *− f* (*µ*_*t−*1_) is the prediction error, and *K*_*t−*1_ is the predictive variance, combining the previous recognition variance with the variance of latent change. The gain *η*_*t*_ controls the direction and strength of the mean update. The mean is updated first, and the precision is then evaluated at the updated mean. Details of the variational formulation and its approximation are described in Konaka and Honda [24].

The source of the observation and the gain depend on the phase. During defeat, each observation is an attack and the gain is *α*:

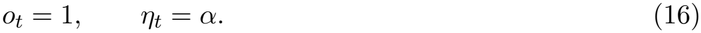

During sensory contact, a latent threat value is sampled from the current recognition distribution, and an observation is generated through the observation model:

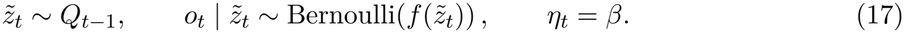

The internally generated observation is then used to update the belief. Repeating this process implements the recursive learning loop. Observations during sensory contact are generated using the current recognition variance 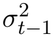, whereas the predictive variance *K*_*t−*1_ enters the recognition update.

The gain *η*_*t*_ scales the mean update, whereas precision follows Eq. (14). During sensory contact, allowing *β <* 0 reverses the direction of the mean update and extends the rule beyond ordinary free-energy minimization. At *β* = 0, the mean remains unchanged, although the precision can still evolve.

### 6.2 Simulation protocol and parameters

Each session included 10 defeat updates, in which *o*_*t*_ = 1 and the gain was *α*, followed by 960 sensory-contact updates, in which *o*_*t*_ was generated from the current recognition distribution according to Eq. (17) and the gain was *β*. The recognition state was carried forward between phases.

All simulations used the initial recognition state 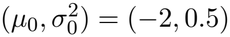 and *v*_*z*_ = 0.1. The simulations in Figs. 2, 3, and 4a,c ran for ten sessions (9,700 updates). For the negative-gain comparison in Fig. 4e, simulations ended after one session (970 updates).

The representative simulations in Figs. 2 and 3 used 1,024 runs per gain at *α* = 0.136, with *β* = 0.48, 1.04, 1.96 and *β* = *−*0.48, *−*1.04, *−*1.96. Random seeds were shared across all six gain conditions.

The parameter survey in Fig. 4a covered 7,171 parameter pairs, with *α* ranging from 0.02 to 0.30 in increments of 0.004 and *β* ranging from *−*2 to 2 in increments of 0.04. The same 20 random seeds were used for every parameter pair.

The distribution comparisons in Fig. 4c,e used 16,384 runs per gain, separate from the runs for Figs. 2 and 3 and the parameter survey. Reported sample SDs quantify variability across simulation runs and use denominator *n −* 1, where *n* is the number of runs.

### 6.3 Mean and variance of belief updates during sensory contact

During sensory contact, the mean threat belief follows Eq. (13) with *η*_*t*_ = *β*. For a fixed pre-update recognition state 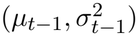, the internally generated observation *o*_*t*_ is stochastic. We calculated the mean and variance of the resulting single-step change,

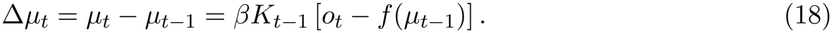

The conditional mean and variance are

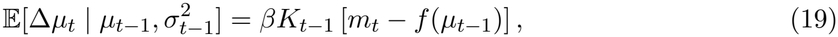

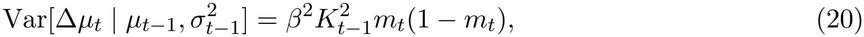

where *m*_*t*_ is the conditional probability of the internally generated attack *o*_*t*_ = 1, averaged over the pre-update recognition distribution:

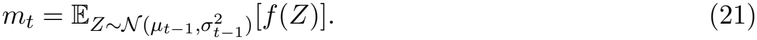

To understand the direction of the mean change, we approximated *m*_*t*_ by expanding *f* (*Z*) to second order around *µ*_*t−*1_. For small recognition variance 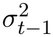,

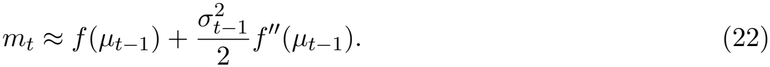

Using *f* ^*′′*^(*u*) = *f* (*u*)[1 *− f* (*u*)][1 *−* 2*f* (*u*)] and 1 *−* 2*f* (*u*) = *−* tanh(*u*/2), the mean change becomes

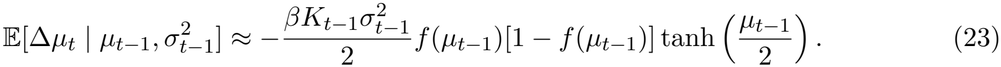

Thus, the mean change is directed toward *µ* = 0 for *β >* 0 and away from it for *β <* 0.

### 6.4 Continuous-time approximation and threat distributions

#### 6.4.1 Continuous-time approximation

To understand how recursive belief updating shapes threat perception during sensory contact, we approximated the discrete updates by a continuous-time stochastic process. Define 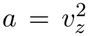 and *h*(*x*) = *f* ^*′*^(*x*) = *f* (*x*)[1 *− f* (*x*)]. For small recognition variance 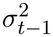, Eq. (22) gives *m*_*t*_(1 *− m*_*t*_) ≃ *h*(*µ*_*t−*1_). Using this approximation in Eq. (20), together with the conditional mean in Eq. (23), yields

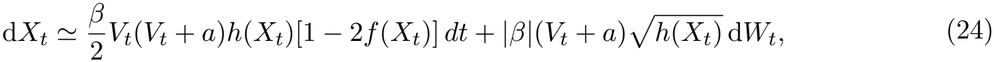

where *X*_*t*_ and *V*_*t*_ are the continuous-time counterparts of *µ*_*t*_ and 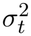, respectively, and *W*_*t*_ is a standard Wiener process.

To obtain an equation for *X*_*t*_ alone, we assumed that *V*_*t*_ follows its local equilibrium value *V*_*_(*X*_*t*_). Holding the belief mean fixed at *X* and setting 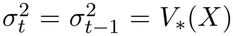 in Eqs. (14) and (15) gives

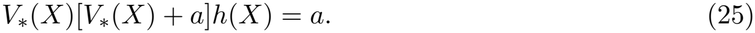

Replacing *V*_*t*_ by *V*_*_(*X*_*t*_) in Eq. (24) and using Eq. (25) gives

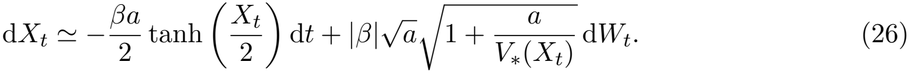

For *a*/*V*_*_(*X*) ≪ 1, the noise amplitude is approximately 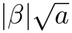, yielding the constant-noise SDE in Eq. (8).

#### 6.4.2 Stationary distribution for positive gains

For the constant-noise SDE in Eq. (8), the probability density *ρ*(*X, t*) of the belief mean *X*_*t*_ satisfies the Fokker–Planck equation

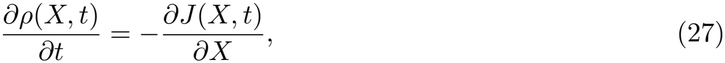

where the probability current *J* (*X, t*) is

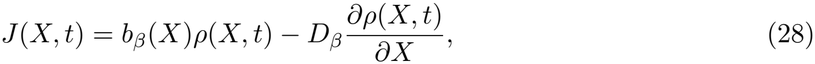

with drift *b*_*β*_ (*X*) and diffusion coefficient *D*_*β*_ given by

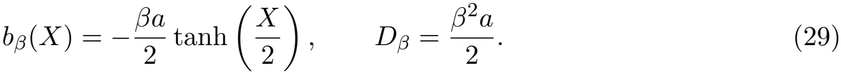

At stationarity, the probability current is constant. For *β >* 0, we consider a normalizable stationary density *ρ*_*β*_(*X*) on the real line and set *J* = 0. Integrating the resulting equation gives

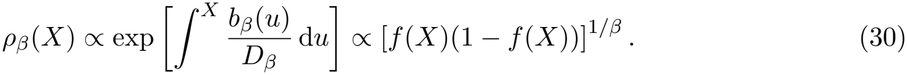

After normalization,

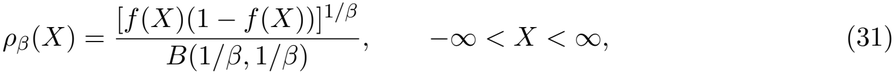

where *B* is the beta function.

Changing variables from *X* to perceived threat *q* = *f* (*X*), the corresponding density is

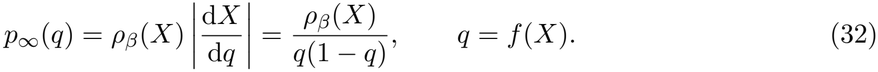

Substituting Eq. (31) gives

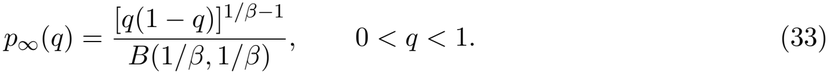

Thus, the stationary distribution of perceived threat is a symmetric beta distribution with shape parameters 1/*β*, denoted Beta(1/*β*, 1/*β*). This distribution is distinct from the beta function *B* used for normalization.

#### 6.4.3 Finite-time distribution for negative gains

For *β <* 0, the Fokker–Planck equation has no normalizable stationary density. We therefore solved the time-dependent equation numerically. Taking *t* = 0 at the start of the first sensory-contact phase, we initialized the density at the belief mean *µ*_D_ reached after ten defeat updates:

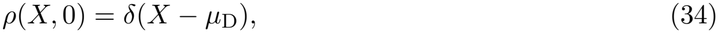

where *δ* is the Dirac delta. Using a finite-volume method with implicit time stepping, we integrated the equation to *t* = 960, corresponding to the end of the first sensory-contact phase (*T* = 970). The resulting distribution of *X*_*t*_ was transformed into the distribution of perceived threat *q* = *f* (*X*_*t*_) for the comparison in Fig. 4e.

## 7 Data availability

This study is purely computational and based entirely on numerical simulations. No experimental or empirical data were generated.

## 8 Code availability

The Python code used for this study is available at https://github.com/word012/rumination-world-model.

## 9 Ethics statement

No animal experiments were performed as part of this computational study.

## 10. Acknowledgements

This study was supported primarily by the Japan Agency for Medical Research and Development (AMED) through the Multidisciplinary Frontier Brain and Neuroscience Discoveries program (Brain/MINDS 2.0, grant 26wm0625322 to T.H., T.A.K., and H.N.). AMED provided additional support through Brain/MINDS 2.0 (grants 26wm0625210 and 26wm0625424 to H.N.) and AMED-CREST (grant 26gm2010011 to H.N.). The Japan Science and Technology Agency (JST) provided support through CREST (grant JPMJCR25Q2 to H.N.), ACT-X (grant JPMJAX26L3 to Y.O.), and BOOST (grant JPMJBS2424 to K.I. and I.H.). The study also received support from Collaborative Research Program of the Institute for Protein Research, the University of Osaka, CR-26-03 (to T.H.), JSPS KAKENHI (grant JP25K24423 to Y.O.) and a Takeda Science Foundation Medical Research Grant (2025046603 to M.T.).

## 11 Author contributions

K.I., M.T., I.H., and H.N. designed the study and developed the model. K.I. and M.T. performed the simulations and analyses. K.I., M.T., Y.O., and H.N. contributed to the mathematical and numerical analyses. T.H. and T.A.K. provided expertise on social defeat stress and contributed to the interpretation of the results. M.T. and H.N. supervised the computational study. K.I., M.T., and H.N. wrote the manuscript and prepared the figures, with input from all authors. All authors reviewed and approved the manuscript.

